# Investigating Carboxysome Morphology Dynamics with a Rotationally Invariant Variational Autoencoder

**DOI:** 10.1101/2021.11.15.468661

**Authors:** Miguel Fuentes-Cabrera, Jonathan K. Sakkos, Daniel C. Ducat, Maxim Ziatdinov

## Abstract

Carboxysomes are a class of bacterial microcompartments that form proteinaceous organelles within the cytoplasm of cyanobacteria and play a central role in photosynthetic metabolism by defining a cellular microenvironment permissive to *CO*_2_ fixation. Critical aspects of the assembly of the carboxysomes remain relatively unknown, especially with regard to the dynamics of this microcompartment. We have recently expressed an exogenous protease as a way of gaining control over endogenous protein levels, including carboxysomal components, in the model cyanobacterium *Synechococcous elongatus* PCC 7942. By utilizing this system, proteins that compose the carboxysome can be tuned in real-time as a method to examine carboxysome dynamics. Yet, analysis of subtle changes in carboxysome morphology with microscopy remains a low-throughput and subjective process. Here we use deep learning techniques, specifically a Rotationally Invariant Variational Autoencoder (rVAE), to analyze the fluorescence microscopy images and quantitatively evaluate how carboxysome shell remodelling impacts trends in the morphology of the microcompartment over time. We find that rVAEs are able to assist in the quantitative evaluation of changes in carboxysome location, shape, and size over time. We propose that rVAEs may be a useful tool to accelerate the analysis of carboxysome assembly and dynamics in response to genetic or environmental perturbation, and may be more generally useful to probe regulatory processes involving a broader array of bacterial microcompartments.

## 1 Introduction

Cyanobacteria are prokaryotic autotrophs that are under investigation as an alternative chassis for the solar-driven conversion of *CO*_2_ into useful bioproducts [1, 2, 3, 4]. Like other members of the green photosynthetic lineage, carbon fixation is accomplished in cyanobacteria through the enzymatic activity of ribulose-1,5-bisphosphate carboxylase/oxygenase (Rubisco) [5]. Among the model cyanobacterial species, *Synechococcus elongatus* PCC 7942 (hereafter *S. elongatus*) has a well-developed genetic toolkit and has been the subject of extensive research on circadian rhythms, carbon metabolism, metabolic engineering, and carboxysome biogenesis.

The carboxysome is a proteinaceous bacterial microcompartment that exists within the cytosol of cyanobacteria, encapsulates a phase-separated pool of Rubisco, and creates a micro-environment favorable to the carboxylation reaction [6, 7]. Despite the carboxysome’s central role in cyanobacterial metabolism, a complete picture of its biogenesis and remodeling remains elusive, though over the years several key studies have provided insights [8, 9, 10]. One central question relates to the degree of dynamic reconfiguration of carboxysomes once assembled. While it is well-documented that carboxysome size, number, distribution, and perhaps shell composition can be modulated to changing environments [11, 12, 13, 14, 15], it is unclear if the observed restructuring is only true for newly-synthesized carboxysomes, or if pre-existing carboxyomes are sufficiently dynamic to be remodeled in response to changing conditions. For example, some evidence suggests that once carboxysomes are formed, they are static until they are ultimately degraded as a unit [10].

Towards a better understanding of carboxysome biogenesis, we sought to develop a quantitative method for investigating the temporal dynamics of carboxysome remodelling. In recent work [16], we used a method for protein down regulation based on the Lon protease from *Mesoplasma florum* (*mf-lon*) allowing rapid inducible degradation of proteins of interest. This approach offers the advantage that carboxysome components can be specifically targeted in a manner that causes dynamic rearrangement of carboxysome morphology beginning at an experimentally defined time point. We previously targeted the carboxysome shell protein CcmO: carboxysomes deficient in CcmO form a distinct phenotype related to their inability to close completely [8]. Using fluorescence microscopy, we previously showed that *mf-lon* degraded CcmO and led to carboxysome remodelling within 24 hours of activation. Here we describe the use of deep learning techniques to analyze the fluorescence microscopy images and quantitatively evaluate trends in carboxysome degradation (Figure 1).

**Figure 1:**
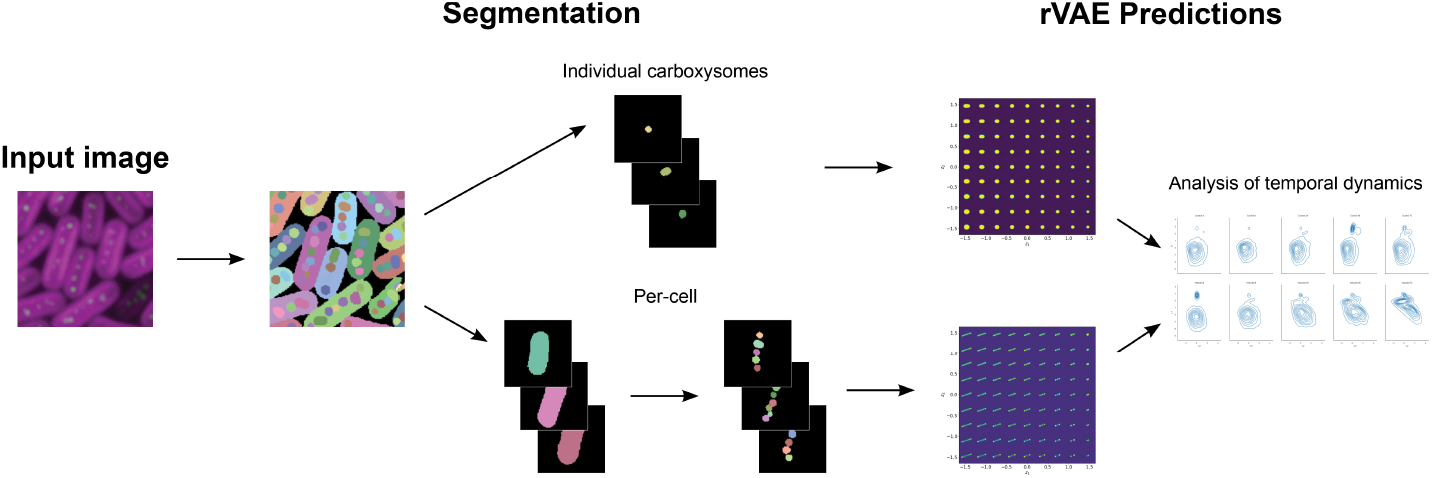
Overview diagram depicting the rVAE workflow

## 2 RESULTS AND DISCUSSION

Elucidating complex biological interactions via microscopy, particularly those which change over time, is a challenging process. Nuanced differences in protein localization, stochasticity, and observational subjectivity often preclude strong conclusions from being made based solely on observation. In this work, we used a deep learning network consisting of a Rotationally-invariant Variational Autoencoder (rVAE) to perform quantitative analysis of carboxysome degradation temporal dynamics.

A detailed explanation of the the segmentation procedure and the image dataset is given in section 4. Here both are summarized for the reader’s convenience. The dataset consisted of images taken at 4, 8, 24, 48 and 72 hours. In order to visualize changes in carboxysome morphology, we tagged Rubisco with a fluorescent protein, mNeonGreen (mNG, see Methods, Section 4). The dataset was divided in two sets, the control and the induced set. Only samples of the latter set contained the exogenous protease that degraded carboxysomes’ shells. Segmentation consists in partitioning an image into its constituents objects. In our case, each image has three channels: brightfield, chlorophyll a autofluorescence (DsRed), and the carboxysomes (mNG). Segmentation was performed for the cell and carboxysome channels using Cellpose [17], a recently developed algorithm specifically trained for cellular segmentation. Figures 2a,b show examples of segmentation for the channels containing cells and carboxysomes, respectively.

**Figure 2:**
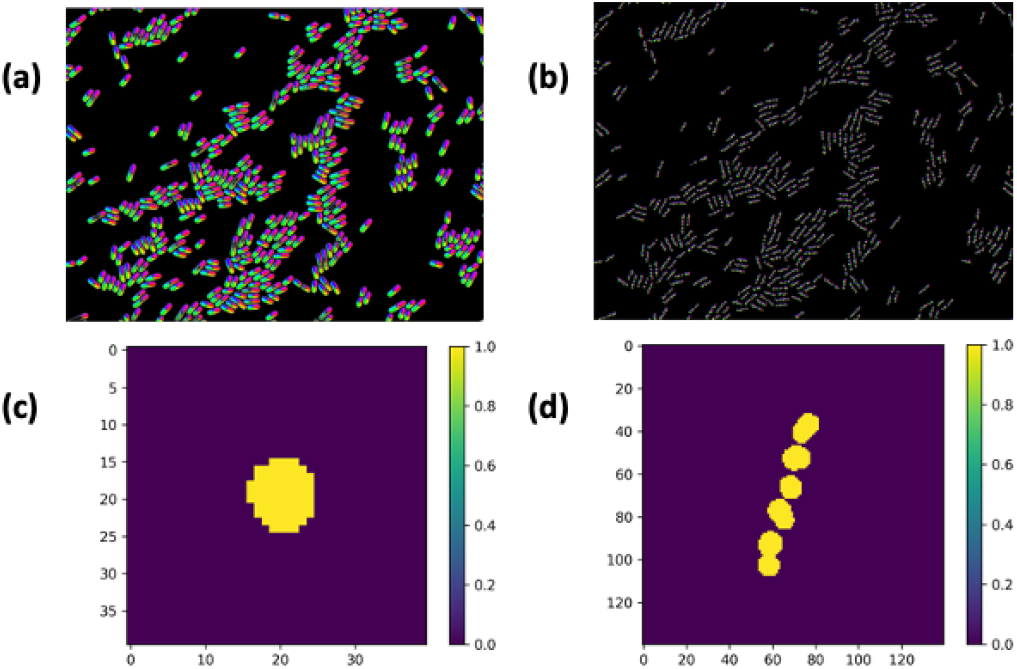
Segmentation of a fluorescence microscopy image shows: (a) the cyanobacteria cells; (b) the carboxysomes within all the cells; (c) an individual carboxysome; (d) the set of carboxysomes within a cyanobacterium.

From the segmented images, we cropped sub-images containing either individual carboxysomes or all the carboxysomes in a cyanobacterium. Examples of cropped sub-images are shown in Figs. 2c,d. We repeated this procedure for each segmented image, generating stacks of sub-images for the control and induced samples. Table 1 summarizes the sub-images stacks and their dimensions. These stacks were analyzed with an rVAE.

**Table 1:**
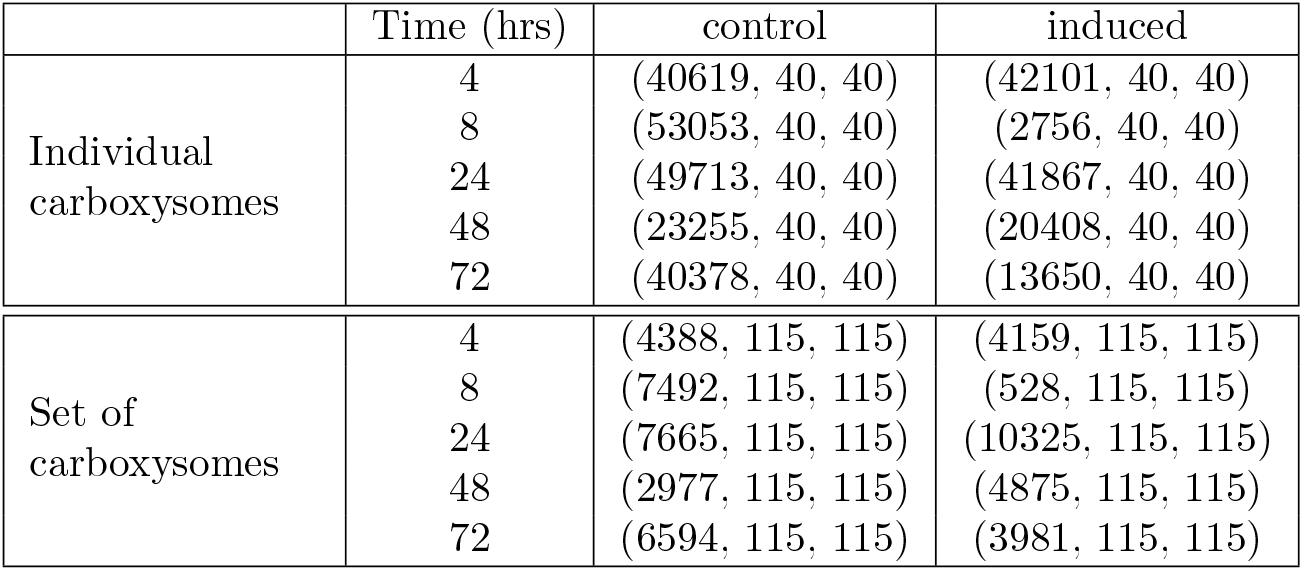
Dimensions of the sub-images stacks for the control and induced samples.

|  | Time (hrs) | control | induced |
| --- | --- | --- | --- |
| Individual<br>carboxysomes | 4 | (40619, 40, 40) | (42101, 40, 40) |
|  | 8 | (53053, 40, 40) | (2756, 40, 40) |
|  | 24 | (49713, 40, 40) | (41867, 40, 40) |
|  | 48 | (23255, 40, 40) | (20408, 40, 40) |
|  | 72 | (40378, 40, 40) | (13650, 40, 40) |
| Set of<br>carboxysomes | 4 | (4388, 115, 115) | (4159, 115, 115) |
|  | 8 | (7492, 115, 115) | (528, 115, 115) |
|  | 24 | (7665, 115, 115) | (10325, 115, 115) |
|  | 48 | (2977, 115, 115) | (4875, 115, 115) |
|  | 72 | (6594, 115, 115) | (3981, 115, 115) |

It is important to explain why we used in our analysis a rVAE and not a “vanilla” Variational Autoencoder, VAE [18]. Both are comprised of an encoder and a decoder network. The encoder compresses images into a low-dimensional representation, known as the latent space. Vectors in this space contain the most relevant information of each image. The decoder reconstructs images starting from the latent space. During encoding, it’s often desirable to disentangle relevant information in the images from mere rotation and translation. Because we wish to understand how degradation affects the structure of the carboxysome, rotations and translations are irrelevant. A rVAE, unlike a VAE, can disentangle rotations and translation during encoding [19], and this is the reason why we used a rVAE, instead of a VAE, in our analysis.

We used the rVAE implementation in Kalinin *et al*. [20] and AtomAI [21]. This rVAE encodes the images into unstructured latent variables, of which we used only two, *L*1 and *L*2, and the latent variables encoding the rotational angle, *L*_*θ*_, and the translation in *x* and *y, L*_Δ*x*_ and *L*_Δ*y*_. Below, we present results for *L*1 and *L*2 only. For *L*_*θ*_, *L*_Δ*x*_ and *L*_Δ*y*_, the results are included in the Supporting Information.

### 2.1 Individual Carboxysomes

Images of individual carboxysomes decoded from the latent space are shown in Fig. 3a, with *L*1 and *L*2 varying along the *x* and *y* axis, respectively. It is seen that carboxysomes can be of different sizes and shapes. For understanding what *L*1 and *L*2 represent, we decoded the images by varying *L*1 and fixing *L*2, and vice versa.

**Figure 3:**
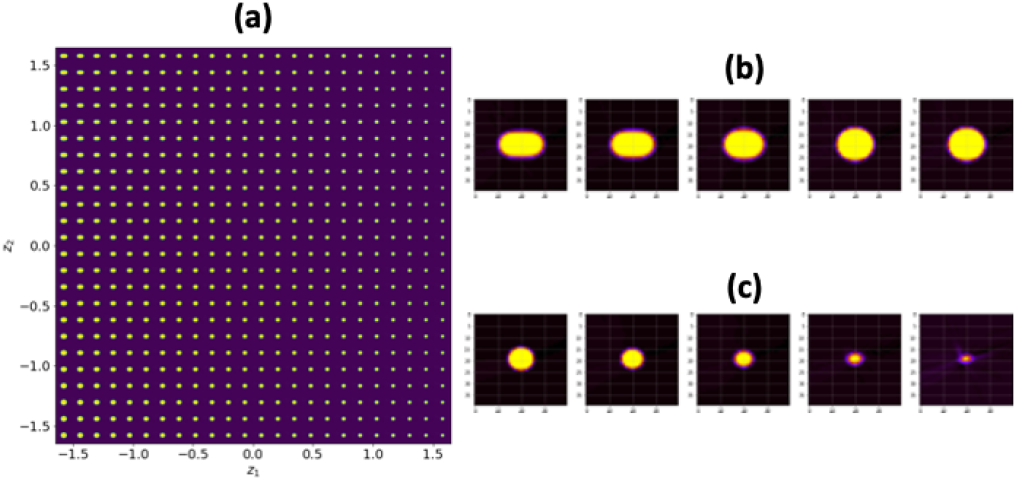
Images of individual carboxysomes decoded from the latent space. (a) Evolution of decoded images as a function of *L*1 and *L*2; (b) decoded images for *L*1 = −1.5, −1, 0, 1, 2 and *L*2 = 2; (c) decoded images for *L*1 = 0 and *L*2 = 0, 1, 2, 3, 4.

The results are shown Figs. 3b,c. It is seen that *L*1 represents the shape of the carboxysome whereas *L*2 represent the size.

The *L*1 and *L*2 histograms for the control and induced samples are shown in Figs. 4 and 5, respectively. The histograms for the control samples remain practically constant with time, whereas the histograms for the induced samples vary. Indeed, in Fig. 4 the *L*1 histograms for the induced samples show a large peak at *L*1 = 0 and a “shoulder” for *L*1 *>* 1.0 that grows steadily with time. In Fig. 5 the *L*2 histograms show a decreasing with time of the large peak at *L*2 = 0 and the appearance of a new peak in the region *L*2 = [−2, −1].

**Figure 4:**
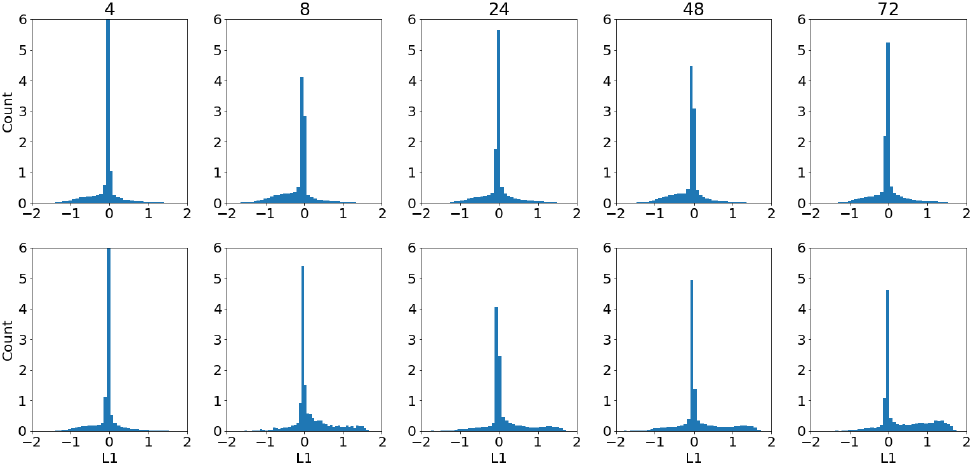
*L*1 histogram for the stacks of individual carboxysomes sub-images, Table 1, for the control and induced samples

**Figure 5:**
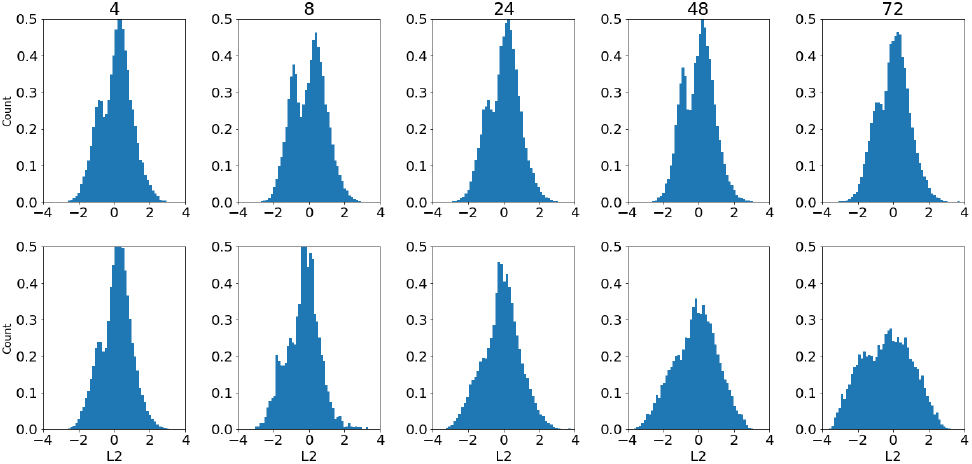
*L*2 histogram for the stacks of individual carboxysomes sub-images, Table 1, for the control and induced samples

The results above indicated that degradation changes *L*1 and *L*2 in the induced samples. To visualize what those changes meant in terms of carboxysomeal structural dynamics, we used the *L*1-*L*2 joint distribution. This distribution is shown in Fig. 6a, and it is seen that at 72 hours there’s an increase in the number of carboxysomes in the regions *L*1 *>* 1 and *L*2 = [−2, −1]. Thus, decoding images from those regions will visualize the structural changes caused by degradation. The corresponding decoded images are shown in Figs. 6b,c,d, where *L*1 increases from bottom to top and *L*2 decreases from left to right. It is clear that degradation produces large round or elongated carboxysomes.

**Figure 6:**
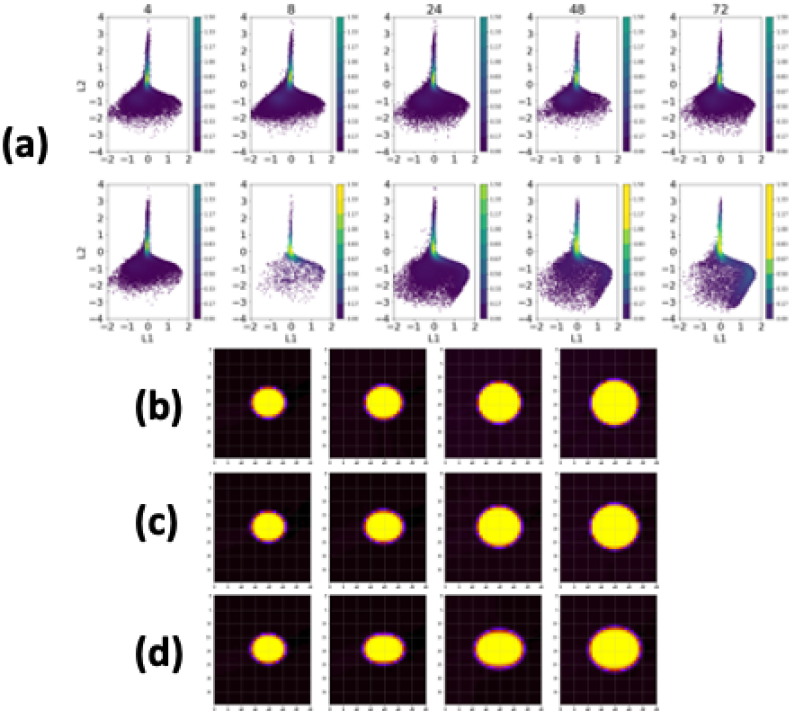
The *L*1-*L*2 joint distribution for the induced samples, with the kernel density estimate superimposed. (a) The *L*1 − *L*2 distribution for each hour; (b)-(d) decoded images from regions of interest (see text).

### 2.2 Set of Carboxysomes per cell

The images of set of carboxysomes decoded from the latent space are shown in Fig. 7a. Figure 7b shows that changing *L*1 while fixing *L*2 generates sets with two carboxysomes located at the poles of the cell. We refer to this effect as “polarization”. Changing *L*2 while fixing *L*1 leads to the collapsing of two carboxysomes into one, which is approximately located at the center of the cell. Both polarization and collapsing decrease the number of carboxysomes per cell.

**Figure 7:**
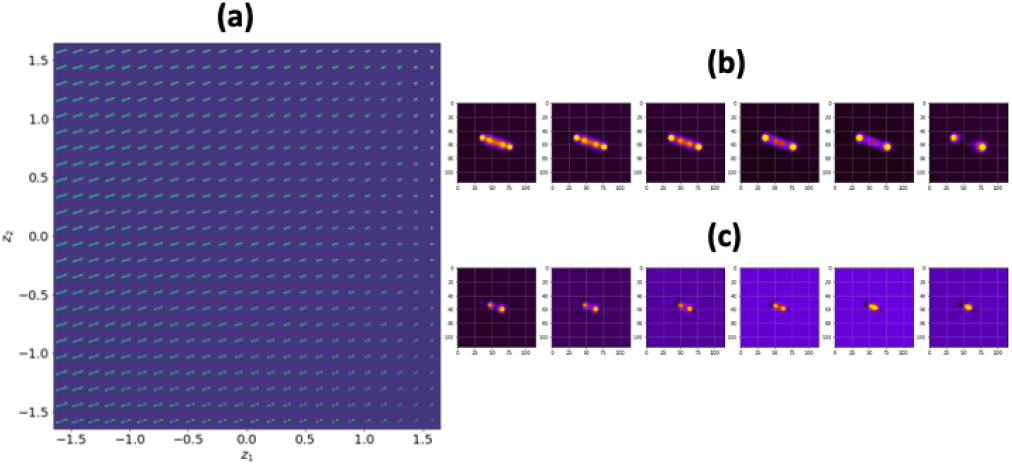
Images of set of carboxysomes decoded from the latent space. (a) Evolution of decoded images as a function of *L*1 and *L*2; decoded images for (b) *L*1 = −2, −1, 0, 1, 1.5 and *L*2 = 0; (c) *L*1 = 1.7 and *L*2 = 1.4, 1.5, 1.6, 1.7, 1.9, 2.

The *L*1, *L*2 histograms for the control and induced samples are shown in Figs. 8, 9, respectively. The *L*1 histogram, Fig. 8, of the control sample remains practically constant with time. For the induced sample, the histogram flattens, widens and moves to larger values of *L*1. Degradation, then, produced more polarization in the induced samples. In contrast, the *L*2 histograms for both the control and induced samples, Fig. 9, change over time. At 48 hours, the *L*2 histogram for the control sample show large peaks at *L*2 *>* 2. For the induced samples, however, large peaks appear as early as at the 4 hour. Additionally, for the induced sample, there is a displacement of the histogram towards larger values of *L*2. Thus, collapsing of carboxysomes happened both in control and induced samples, but it’s more pronounced in the latter.

**Figure 8:**
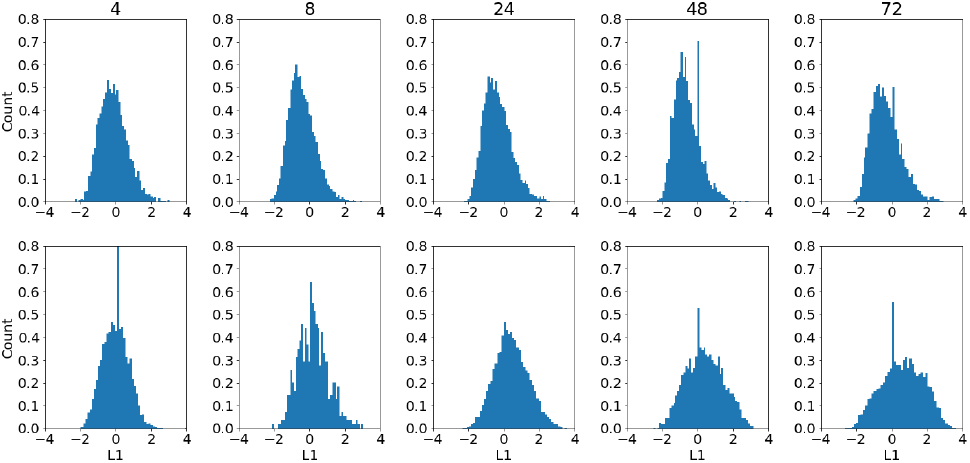
*L*1 histogram for the set of carboxysomes’ sub-images, Table 1, for the control and induced samples.

**Figure 9:**
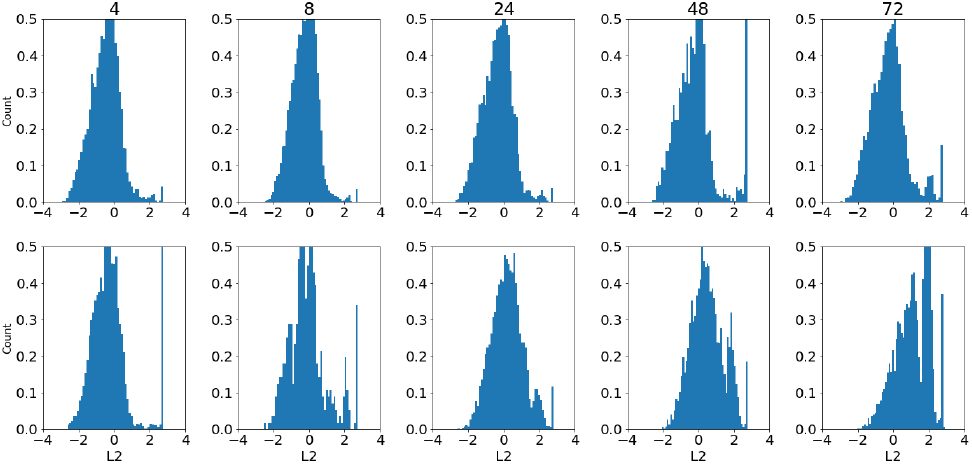
*L*2 histogram for the set of carboxysomes sub-images, Table 1, for the control and induced samples.

The extend to which collapsing and polarization is more significant in the induced samples can be seen in the *L*1-*L*2 joint distribution, Fig. 10. For the control samples, there is an increase in *L*2 *>* 2 values starting at the 4 hour. For the induced samples, the increase in the *L*2 *>* 2 values also occurs early on, but it’s significantly more dramatic, to the point that it ends up splitting the joint distribution into two regions below and above *L*2 = 2. For the control samples, the *L*1 values practically did not changed, whereas for the induced samples *L*1 values changed so much that the distribution is skewed to the right.

**Figure 10:**
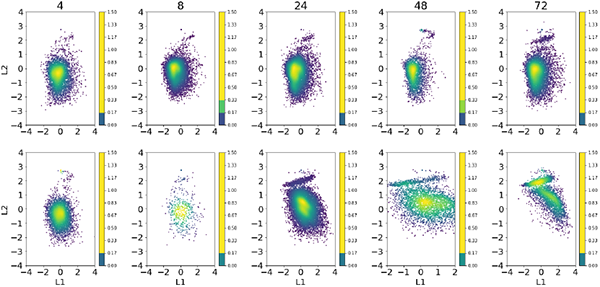
*L*1 − *L*2 joint distributions for the sub-images of the set of carboxysomes for the control and induced samples.

Figure 11 shows decoded images in regions of interest in the *L*1 − *L*2 joint distribution at the 72 hour. Figures 11a,b contain images decoded in the “skewed to the right” region of the joint distribution. The results reveal how degradation clearly increases polarization. Figures 11c,d contain images decoded in the region above *L*2 = 2, and it is clear that collapsing increases also with degradation.

**Figure 11:**
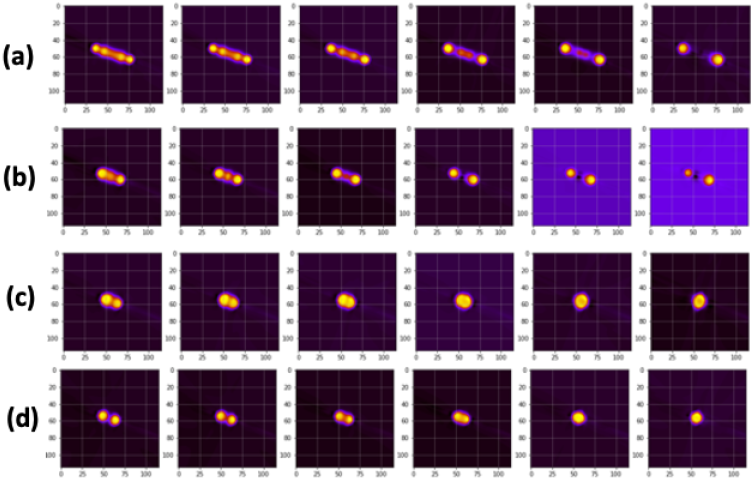
Decoded images from the *L*1 − *L*2 joint distributions for the the selected regions. (a) *L*1 = −2, −1, 0, 1, 1, 5, 2.5, *L*2 = 0; (b) *L*1 = −2, −1, 0, 1, 1, 5, 2.5, *L*2 = 1; (c)*L*1 = 0, *L*2 = 1.4, 1.5, 1.6, 1.7, 1.9, 2.0; (d) *L*1 = −1.5, *L*2 = 1.4, 1.5, 1.6, 1.7, 1.9, 2.0

## 3 CONCLUSIONS

Investigating the biogenesis of carboxysomes can have important implications for the production of commodity chemicals and high-valued products. In this context, imaging with fluorescence microscopy while disturbing the carboxysome can provide valuable insights on structure dynamics. However, as imaging techniques become more efficient, the amount of data produced is such that manual evaluation is so costly that only a qualitative analysis is possible. Quantitative analysis can shed more light into the carboxysome’s biogenesis, and for this purpose, deep learning techniques are especially suitable. Here the demonstrate that a type of deep learning technique known as a Rotationally Invariant Variational Autoencoder is capable of revealing structural changes that happen during degradation, from changes in the shape and size of carboxysomes, to spatial rearrangements inside of the cell that include polarization and collapsing. This work reveals that variational autoencoders can play a very important role in understanding the formation of not only carboxysomes but bacterial micro-compartments in general.

## 4 METHODS

### 4.1 Microbial culturing conditions

*S. elongatus* cultures were grown in baffled flasks (Corning) with BG-11 medium (Sigma) supplemented with 1 g/L HEPES, pH 8.3, in a Multitron II shaking incubator (Infors HT). Cultures were grown under continuous light with GroLux bulbs (Sylvania) at 125 *μ* mol photons *m*^−2^*s*^−1^, 2 % *CO*_2_, 32 °C, and 130 rpm shaking. Carboxysomes were degraded by inserting a protein degradation tag (PDT) at the C-terminus of the shell protein CcmO, which is essential for carboxysome closure, and expression of a non-native Lon protease from *Mesoplasma florum* [16]. For visualization of changes in carboxysome morphology, a second copy of the small subunit of Rubisco, rbcS, was tagged with a C-terminal fusion of mNeonGreen (mNG). For the induced samples, cultures were induced with 30 *μM* theophylline. More detailed culturing, genetic assembly, and transformation information was previously described [16].

### 4.2 Microscopy

All experiments were performed on live cells in exponential growth. Images were collected with a Zeiss Axio Observer D1 inverted microscope with a Zeiss Plan Apochromat 100× lens. Epifluorescence images were collected of both chlorophyll *a* autofluorescence (*λ*_*ex*_ = 545, *λ*_*em*_ = 605) and mNG (*λ*_*ex*_ = 500, *λ*_*em*_ = 535).

### 4.3 Segmentation

A total of 90 images, each with dimensions (1460,1936) pixels, were segmented with Cellpose [17] and analyzed with an rVAE. The set of 90 images is divided into control and induced sample groups, which contain 42 and 39 images, respectively. Both groups contain a time series with images collected at 4, 8, 24, 48, and 72 hours post induction.

Segmentation was performed with Cellpose [17], and for each image, masks for the carboxysomes and the cells were created. We used a diameter setting of 7 and 20 for generating the masks for carboxysomes and the cells, respectively.

From these masks, two types of sub-image stacks were obtained. One stack contains only individual carboxysomes, whereas the other contains the set of carboxysomes per cell. The latter was generated by using the cell channel mask, within which we selected the set of carboxysomes in each cell. The resultant images were padded to have a size of 115 × 115 pixels. The dimension of each stack for the control and induced groups are given in Table 1.

As it can be seen in Table 1, there are significantly less sub-images in the stacks for the set of carboxysomes than in the stack for individual ones. This is due to the position of the focal plane, which is inclined in some of the samples and causes some group of cells to be visualized different than others. This affects the segmentation, and the end result is that there are fewer cell masks than expected. Because these masks are used as the molds within which we selected the set of carboxysomes sub-images, the stacks for the set of carboxsyomes have less sub-images than the stacks for the individual ones.

### 4.4 rVAE analysis

The rVAE was implemented within the AtomAI Package [21] and trained on the subimages stack for 1000 iterations using 3-layer perceptron for the encoder and decoder. Each layer had 128 neurons and was activated by the *tanh*() function, whose weights were optimized using the Adam optimizer with a learning rate of 0.0001.

## ACKNOWLEDGMENTS

This research was conducted at the Center for Nanophase Materials Sciences, which is a DOE Office of Science User Facility. This work was also funded by National Science Foundation Grant 1517241 (to D.C.D.), funding from Department of Energy Grant *DE* − *FG*02 − 91*ER*20021 (to D.C.D.) at the MSU DOE-PRL. Additional support for the research was provided by NSF Award 1845463 (to D.C.D.).

## Licenses and Permissions

This manuscript has been authored by UT-Battelle, LLC under Contract No. DEAC05-00OR22725 with the U.S. Department of Energy. The United States Government retains and the publisher, by accepting the article for publication, acknowledges that the United States Government retains a non-exclusive, paid-up, irrevocable, world-wide license to publish or reproduce the published form of this manuscript, or allow others to do so, for United States Government purposes. The Department of Energy will provide public access to these results of federally sponsored research in accordance with the DOE Public Access Plan (http://energy.gov/downloads/doe-public-access-plan).

